# Immunological tolerance, pregnancy and pre-eclampsia: the roles of semen microbes and the father

**DOI:** 10.1101/198796

**Authors:** Louise C. Kenny, Douglas B. Kell

## Abstract

Although it is widely recognised as involving two stages (poor placentation followed by oxidative stress/inflammation), the precise originating causes of pre-eclampsia (PE) remain elusive. We have previously brought together some of the considerable evidence that a (dormant) microbial component is commonly a significant part of its aetiology. However, apart from recognising, consistent with this view, that the many inflammatory markers of PE are also increased in infection, we had little to say about immunity, whether innate or adaptive. In addition, we focussed on the gut, oral and female urinary tract microbiomes as the main sources of the infection. We here marshall further evidence for an infectious component in PE, focussing on the immunological tolerance characteristic of pregnancy, and the well-established fact that increased exposure to the father’s semen assists this immunological tolerance. As well as these benefits, however, semen is not sterile, microbial tolerance mechanisms may exist, and we also review the evidence that semen may be responsible for inoculating the developing conceptus with microbes, not all of which are benign. It is suggested that when they are not, this may be a significant cause of preeclampsia. A variety of epidemiological and other evidence is entirely consistent with this, not least correlations between semen infection, infertility and PE. Our view also leads to a series of other, testable predictions. Overall, we argue for a significant paternal role in the development of PE through microbial infection of the mother via insemination.

> “In one of the last articles which he wrote, the late Professor F J Browne (1958) expressed the opinion that all the essential facts about pregnancy toxaemia are now available and that all that is required to solve the problem is to fit them together in the right order, like the pieces of a jigsaw puzzle” [1]

> “It appears astonishing how little attention has been given in reproductive medicine to the maternal immune system over the last few decades.” [2]

## Introduction

Pre-eclampsia (PE) is a multifactorial disease of pregnancy, in which the chief manifestations are hypertension and proteinuria [3-11]. It affects some 3-5% of nulliparous pregnancies worldwide [10; 12; 13], and is associated (if untreated) with high morbidity and mortality [14-18]. There is much literature on accompanying features, and, notwithstanding possible disease subdivisions [19; 20], the development of PE is typically seen as a ‘two-stage’ process (e.g. [21-27]), in which in a first stage incomplete remodelling of spiral arteries leads to poor placentation. In a second stage, the resulting stress, especially hypoxia-induced oxidative stress [28] (and possibly hypoxia-reperfusion injury), then leads to the symptoms typical of later-pregnancy pre-eclampsia. However, the various actual originating causes of either of these two stages remain obscure. Many theories have been proposed (albeit a unitary explanation is unlikely [19]), and indeed, PE has been referred to as a ‘disease of theories’ [1; 29; 30]. The only effective ‘cure’ is delivery [31], which often occurs significantly pre-term, with its attendant complications for both the neonate and in later life. Consequently, it would be highly desirable to improve our understanding of the ultimate causes of PE, so that better prevention or treatments might be possible.

The ‘two-stage’ theory is well established, and nothing we have to say changes it. However, none of this serves to explain what ‘initiating’ or ‘external’ factors are typically responsible for the poor placentation, inflammation, and other observable features of PE [32].

Microbes are ubiquitous in the environment, and one potential external or initiating factor is low-level microbial infection. In a recent review [32], we developed the idea (and summarised extensive evidence for it) that a significant contributor to pre-eclampsia might be a (largely dormant [33-36] and non-replicating) microbiome within the placenta and related tissues, also detectable in blood and urine. Others (e.g. [37-44]) have drawn similar conclusions. Interestingly, recent analyses [19; 45] of placental gene expression in PE implicate changes in the expression of TREM1 (triggering receptor on myeloid cells-1) and the metalloprotease INHA, and in one case [19] also LTF (lactotransferrin), that also occur during infection [46-49]. Although we highlighted the role of antibiotics as potentially preventative of PE [32], and summarised the significant evidence for that, we had relatively little to say about immunology, and ignored another well-known antidote to infectious organisms in the form of vaccines. There is certainly also an immune component to preeclampsia (e.g. [24; 50-58] and below). One of the main theories of (at least part of the explanation of) PE is that of ‘immune maladaptation’ [50-52; 59]. Thus, the main focus of the present analysis is to assess the extent to which there is any immunological evidence for a role of infectious agents (and the utility of immunotolerance to or immunosuppression of them) in PE. Figure 1 summarises our review in the form of a ‘mind map’ [60]. We begin with the broad question of immunotolerance, before turning to an epidemiological analysis.

**Figure 1.**
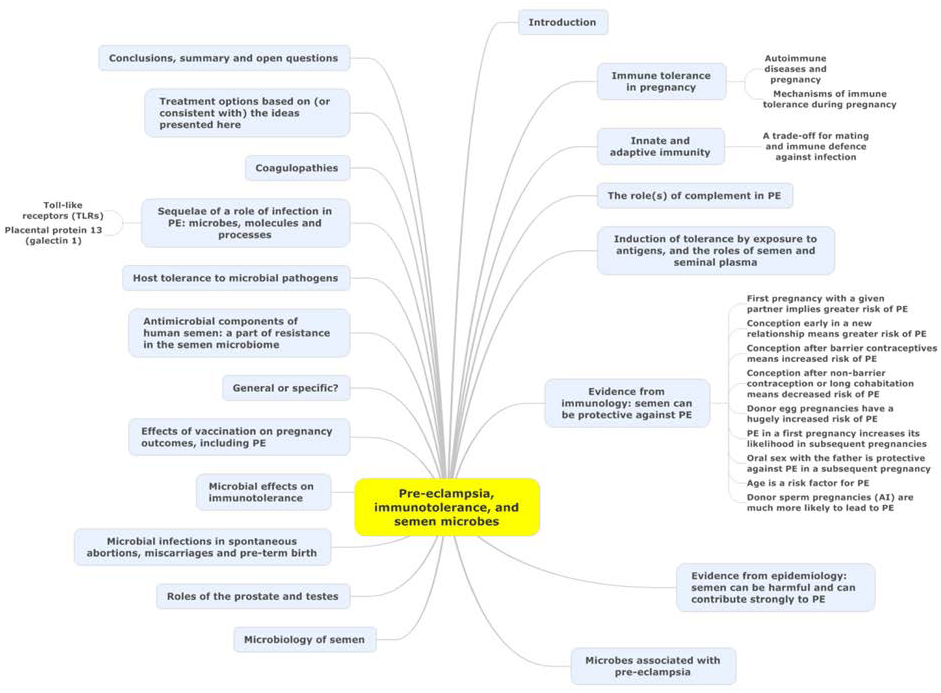
A ‘mind map’ [60] of the review. Start at ‘midnight’ and read clockwise.

## Immune tolerance in pregnancy

Much of the original thinking on this dates back to Sir Peter Medawar [61-66], who recognised that the paternal origin of potentially half the antigens of the fetus [67] created an immunological conundrum: it should normally be expected that the fetus’s alloantigens would cause it to be attacked by the maternal immune system as ‘foreign’. There would therefore have to be an ‘immune tolerance’ [65; 68-70]. Historically it was believed that the fetus is largely ‘walled off’ from the mother [71]; however, we now appreciate [72; 73] that significant trafficking of fetal material across the placenta into the maternal circulation and vice versa occurs throughout pregnancy. Indeed, this is the basis for the development of non-invasive prenatal testing (NIPT). In line with this, grams of trophoblast alloantigens are secreted daily into the maternal circulation during the third trimester (Figure 2), and this is related to the prevalence of PE [74-80]. Consequently, both the concept and the issue of immune tolerance are certainly both real and important. At all events, the immunobiology of the fetus has been treated in theory largely in the way that a transplanted graft is treated, and uteroplacental dysfunction (leading to PET and IUGR) is largely regarded as a graft rejection (e.g. [53; 81-87]). Clearly there are relationships between the immunogenicity of the foreign agent and the responsiveness of the host; to this end, Zelante and colleagues [88] recognise the interesting similarities between tolerance to paternal alloantigens (as in pregnancy) and the tolerance observed in chronic fungal infections.

**Figure 2.**
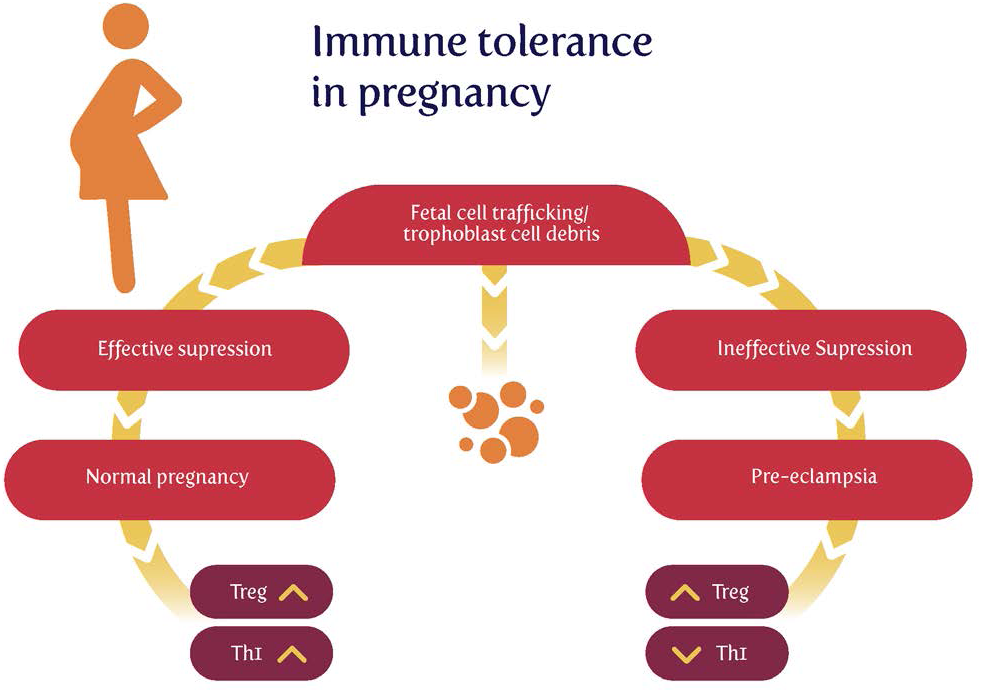
Effective suppression of response to fetal cell trafficking leads to a normal pregnancy, while its failure can lead to pre-eclampsia.

### The clinical course of automimmune disease during pregnancy: an inconsistent effect

The seminal observation by Philip Hench that the symptoms of the rheumatoid arthritis (RA) were frequently and dramatically ameliorated by several conditions, including pregnancy [89], led to the discovery of cortisone [90] and gave unique insights into the complex interaction between the maternal immune system and the developing fetal/placental unit. Contemporary data suggests that the improvement in RA is not ubiquitous as first thought. Amongst all pregnant women about 25% of women have no improvement in their symptoms at any stage in pregnancy and in a small number of cases the disease may actually worsen [91]. The process by which pregnancy affects disease activity in RA is not completely understood and several putative mechanisms have been proposed. Of interest, although plasma cortisol rises during pregnancy and was initially thought to be key in the amelioration of symptoms, there is actually no correlation between cortisol concentrations and disease state [92]. It has also been reported that the degree of maternal and paternal MHC mismatch has been shown to correlate with the effect of the RA remission during pregnancy [93], leading to the hypothesis that the early immunological events in pregnancy that establish tolerance to the fetal allograft contribute to RA remission. Clearly, this may also account for the disparity in response to pregnancy. RA is not unique in being the only autoimmune disease to be profoundly altered by pregnancy. Although less well studied, non-infectious uveitis tends to improve during pregnancy from the second trimester onwards, with the third trimester being associated with the lowest disease activity [94]. Again, the mechanism underlying this phenomenon is not completely elucidated.

It is now generally accepted [95] that, notwithstanding the sweeping generalisation, autoimmune diseases with a strong cellular (innate) pathophysiology (RA, Multiple Sclerosis (MS)) improve, whereas diseases characterised by autoantibody production such as systemic lupus erythematous (SLE) and Grave’s disease tend towards increased severity in pregnancy.

We have previously reported an association between pregnancy and the risk of subsequent maternal autoimmune disease which was also related to the mode and gestation of delivery. There was an increased risk of autoimmune disease after Caesarean section may be explained by amplified fetal cell traffic at delivery, while decreased risks after abortion may be due to the transfer of more primitive fetal stem cells [96].

### Mechanisms of immune tolerance during pregnancy

Following the recognition of maternal immunotolerance, a chief discovery was the choice of HLA-G, a gene with few alleles, for the antigens used at the placental interface. Thus, the idea that placental HLA-G proteins facilitate semiallogeneic pregnancy by inhibiting maternal immune responses to foreign (paternal) antigens via these actions on immune cells is now well established [97-102].

It is also well established that Regulatory T cells (Tregs) play an indispensable role in maintaining immunological unresponsiveness to self-antigens and in suppressing excessive immune responses deleterious to the host [103]. Consequently, much of present thinking seems to involve a crucial role for regulatory Tregs in maintaining immunological tolerance during pregnancy [53; 64; 104-114], with the result that effector T cells cannot accumulate within the decidua (the specialized stromal tissue encapsulating the fetus and placenta) [115].

In an excellent review, Williams and colleagues [116] remark “Regulatory T cells (Tregs) are a subset of inhibitory CD4+ helper T cells that function to curb the immune response to infection, inflammation, and autoimmunity”. “There are two developmental pathways of Tregs: thymic (tTreg) and extrathymic or peripheral (pTreg). tTregs appear to suppress autoimmunity, whereas pTregs may restrain immune responses to foreign antigens, such as those from diet, commensal bacteria, and allergens”. Their differential production is controlled by a transcription factor called Foxp3.

Further, “a *Foxp3* enhancer, conserved noncoding sequence 1 (CNS1), essential for pTreg but dispensable for tTreg cell generation, is present only in placental mammals. It is suggested that during evolution, a CNS1-dependent mechanism of extrathymic differentiation of Treg cells emerged in placental animals to enforce maternal-fetal tolerance [117]”.

Williams and colleagues conclude that “These findings indicate that maternal – fetal tolerance to paternal alloantigens is an active process in which pTregs specifically respond to paternal antigens to induce tolerance. Thus, therapies should aim not to suppress the maternal immune system but rather to enhance tolerance. These findings are consistent with an increase in the percentage of Tregs during pregnancy and with no such increase in women with recurrent pregnancy loss [118]” [116]. Thus maternal tolerance is based on exposure to the paternal alloantigens, although mechanisms such as the haem oxygenase detoxification of haem from degrading erythrocytes [119] are also important. Note too that pregnancy loss is often caused by automimmune activity [120] (and see later).

Additionally, Treg cells have several important roles in the control of infection (e.g. [121-126]). These include moderating the otherwise potentially dangerous response to infection, and being exploited by certain parasites to induce immunotolerance.

Finally, here, it is also recognised that the placenta does allow maternal IgG antibodies to pass to the fetus to protect it against infections. Also, foreign fetal cells persist in the maternal circulation [127] (as does fetal DNA, nowadays used in prenatal diagnosis). One cause of pre-eclampsia is clearly an abnormal immune response towards the placenta. There is substantial evidence for exposure to partner’s semen as prevention for pre-eclampsia, largely due to the absorption of several immune modulating factors present in seminal fluid [128]. We discuss this in detail below.

## Innate and adaptive immunity

Although they are not entirely independent [129; 130], and both respond to infection, it is usual to discriminate (the faster) innate and (the more leisurely) adaptive immune responses (e.g. [131-135]). As is well known (reviewed recently [136]), the innate immune system is responsible for the recognition of foreign organisms such as microbes. It would be particularly convenient if something in the immune response did actually indicate an infection rather than simply any alloantigen, but unfortunately – especially because of the lengthy timescale over which PE develops – innate responses tend to morph into adaptive ones. This means (i) that there may be specific signals from <u>early</u> innate events that may be more or less specific to innate responses, and (ii) that it also does not exclude the use of particular <u>patterns</u> of immune responsive elements [137-139] to characterise disease states.

A dysregulation of the immune system is widely recognised as an accompaniment to normal pregnancy [64; 111; 140-142], and especially in PE [51; 53; 54; 56-59; 143-150], and it is worth looking at it a little more closely.

The innate immune system responds to microbial components such as LPS via cell membrane receptors. Innate immune cells express a series of evolutionarily conserved receptors known as pattern-recognition receptors (PRRs). PRRs recognise and bind conserved sequences known as pathogen-associated molecular patterns (PAMPs). Bacterial lipopolysaccharide (LPS) and peptidoglycan, and double stranded viral RNA are unique to microbes and act as canonical PAMPs, while the main family of PRRs is represented by the Toll-like receptors (TLRs) [151; 152]. Downstream events, as with many others [153; 154] converge on the NF-κB system and/or interferon, leading to the release of a series of inflammatory cytokines such as IL-2, IL-6, IL-8, TNF-α and especially IL-1β.

Matzinger ‘s “danger model” [155-160] (and see [65] and Figure 3) suggested that activation of the immune system could be evoked by danger signals from endogenous molecules expelled from injured/damaged tissues, rather than simply from the recognition of non-self (although of course in the case of pregnancy some of these antigens are paternal alloantigens). Such endogenous molecules are referred to as Damage-associated molecular patterns (DAMPs), but are not our focus here, albeit they likely have a role in at least some elements of PE [161]. We shall see later, however, that Matzinger’s theory is entirely consistent with the kinds of microbial (and disease) tolerance that do seem to be an important part of pregnancy and PE (and see [162]).

**Figure 3.**
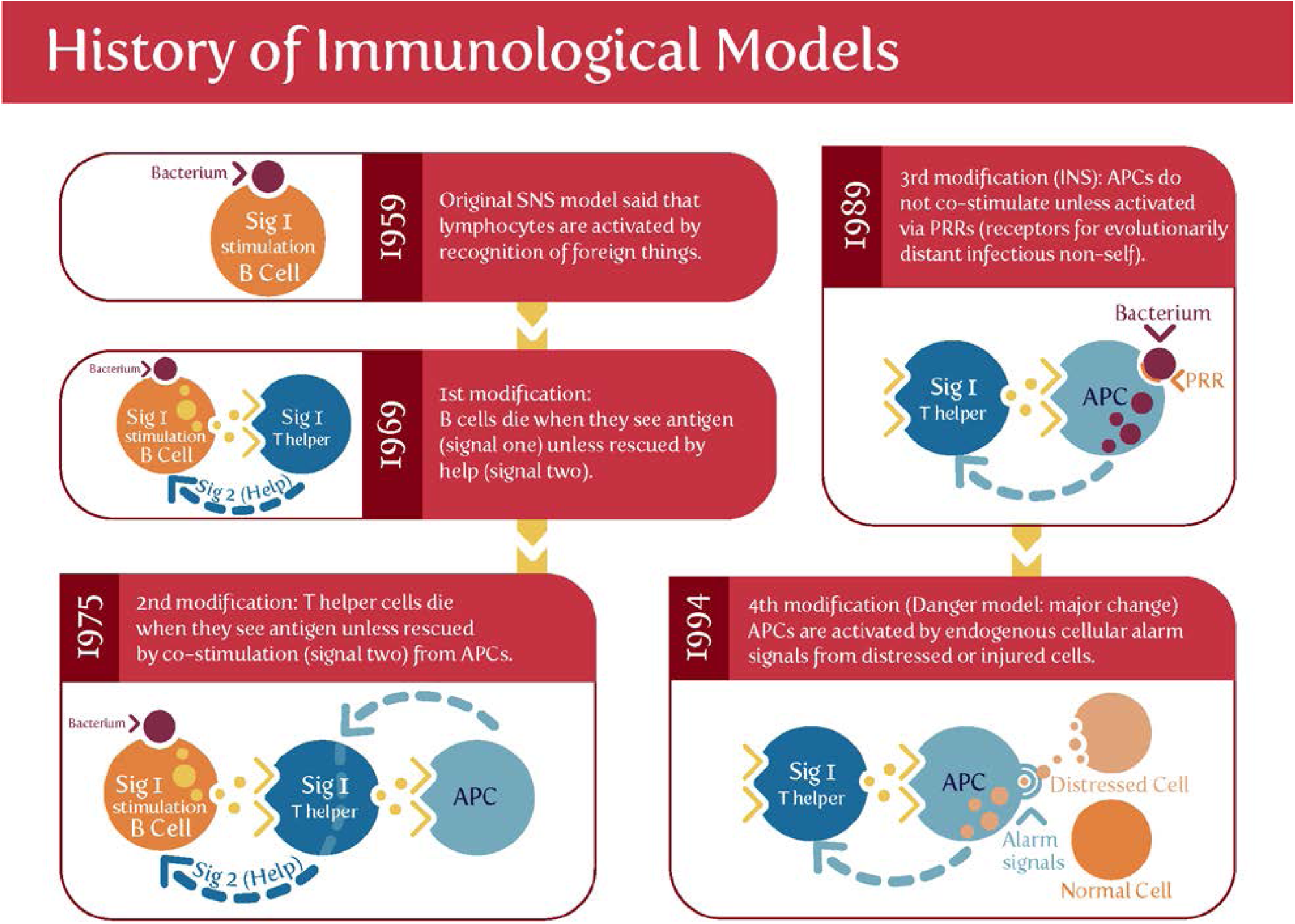
Matzinger’s ‘danger model’ *vs* the classical theory of self vs self-nonself. Based on and redrawn from [158].

The maternal innate immune system plays an important role both in normal pregnancy, and in particular in hypertensive disorders of pregnancy including preeclampsia (PE) [143; 163-169]. One persuasive and widely accepted view is that normal pregnancy is characterised by a low-grade systemic inflammatory response and specific metabolic changes, and that virtually all of the features of normal pregnancy are simply exaggerated in pre-eclampsia [32; 163; 170; 171]. Certainly it is long established that “Normal pregnancy and preeclampsia both produce inflammatory changes in peripheral blood leukocytes akin to those of sepsis” [163], and there are innate Immune defences in the uterus during pregnancy [140]. Normal pregnancy is considered to be a Th2 type immunological state that favours immune tolerance in order to prevent fetal rejection [119]. By contrast, preeclampsia (PE) has been classically described as a Th1/Th2 imbalance [106; 145; 172-174], but as mentioned above (and before [32]), recent studies have highlighted the role of regulatory T-cells as part of a Th1/Th2/Th17 paradigm [143; 144]. This leads to the question of whether there is some kind of trade-off between the responses to paternal alloantigens and those of microbes.

### A trade-off for mating and immune defence against infection

Certainly there is some evidence for a trade-off between mating and immune defence against infection [175-177]. Consistent with this (albeit with much else) is the fact [178-180] that pregnancy is associated with an increased severity of at least some infectious diseases. There is evidence [181; 182] that “adaptive immune responses are weakened, potentially explaining reduced viral clearance. Evidence also suggests a boosted innate response, which may represent a compensatory immune mechanism to protect the pregnant mother and the fetus and which may imply decreased susceptibility to initial infection [179]”.

## The role(s) of complement in PE

Complement, or more accurately the complement cascade, is an important part of the innate immune system that responds to infection. Later (downstream) elements also respond to the adaptive immune system. Our previous review [32] listed many proteins whose concentrations are changed in both infection and PE. Since we regard low-level infection as a major cause of the inflammation observed in PE, one would predict that the complement system is activated in PE, and this observation is amply borne out [183-198]. We give some of the details in Table 1.

**Table 1.** Changes in the Complement system during PE and related pregnancy disorders

| Complement element | Details | Reference |
| --- | --- | --- |
| Bb | Raised in PE, OR 2.1 (CI = 1.4–3.1, P < 0.0003) | [185] |
| Bb | Adjustment for risk factors did not attenuate the association between an elevated Bb and preeclampsia (adjusted odds ratio OR 3.8, 95% CI, 1.6 to 9, P<0.002) in the cohort. After removing women with plasma obtained before 10 weeks, the adjusted OR of Bb in the | [183] |
| | top decile for preeclampsia was 6.1 (95% CI 2.2 to 17, $P < 0.0005$) | |
| Bb | Median Bb levels were higher in the maternal plasma of severe PE subjects ($n = 24$) than in controls ($n = 20$), $1.45 \pm 1.03$ versus $0.65 \pm 0.23 \mu\text{g.mL}^{-1}$, $P < 0.001$. | [196] |
| Bb | Pre-term birth. Women with Bb in the top quartile were 4.7 times more likely to have an SPTB less than 34 weeks' gestation as compared with women who had levels of Bb in the lower 3 quartiles (CI 1.5-14, $P < 0.003$). | [184] |
| Bb | Maternal Bb levels were significantly higher in the preeclamptic group than in the nonpreeclamptic group ($P < 0.003$ in all studied, $P < 0.007$ in African Americans). | [205] |
| Bb | Pyelonephritis. Pregnant women with pyelonephritis had a higher median plasma concentration of fragment Bb than those with a normal pregnancy (1.3 mg/ml, IQR: 1.1–1.9 vs. 0.8 mg/ml, IQR: 0.7–0.9; $P < 0.001$). No significant differences were observed in the median maternal plasma concentration of fragment Bb between pregnant women with pyelonephritis who had a positive blood culture and those with a negative blood culture | [206] |
| Bb | Median amniotic fluid Bb levels were also significantly higher ($P = 0.03$) in preeclamptic women than in normal pregnant women (1127 ng/mL versus 749 ng/mL). The alternative complement pathway is principally involved | [194] |
| Bb, C3a, C5a, and MAC | Increased significantly in EOSPE (all $P < 0.01$) and LOSPE ($P$ value: 0.027, $< 0.001$, 0.001, and $< 0.001$, respectively) compared with Early/Late control. | [195]. See also [197] |
| Bb or C3a | Women who were obese with levels of Bb or C3a in the top quartile were 10.0 (95% confidence interval, 3.3–30) and 8.8 (95% confidence interval, 3–24) times, respectively, more likely to develop preeclampsia compared with the referent group at 20 weeks gestation | [207] |
| C1q and C4d | Increased significantly in LOSPE ($P$ value: .003 and .014, respectively) compared with L-control | [195]. See also [197] |
| C3a | Adjusted for parity and prepregnancy body mass index, women with levels of C3a in the upper quartile in early pregnancy were three times more likely to have an adverse outcome later in pregnancy compared with women in the lowest quartile (95% confidence interval, 1.8–4.8; $P < .001$). This was especially the case for pre-term birth ($P < .0004$). Elevated C3a as early as the first trimester of pregnancy is an independent predictive factor for adverse pregnancy outcomes, suggesting that complement-related inflammatory events in pregnancy contribute to the subsequent development of poor outcomes at later stages of pregnancy | [188] |
| C3a | Autoantibody-mediated complement C3a receptor activation contributes to the pathogenesis of preeclampsia | [191] |
| C3a | Women who developed early-onset preeclampsia as compared with the term pregnant controls had significantly higher ($P = 0.04$) median amniotic fluid C3a levels (318.7 ng/mL versus 254.5 ng/mL) | [194] |
| C3a | 751.6 (194.6–1660) vs 1358 (854.8–2142) $\text{ng.mL}^{-1}$, $P < 0.05$ pre-eclamptic vs healthy pregnant. | [208] |
| C3a, C3a_desArg & C5a | Elevated at term in PE but not earlier (P<0.05) | [209; 210] |
| C3a, C5a & AT1-AA | Levels in serum from the severe pre-eclampsia group were significantly higher than in controls (p < 0.05). | [211] |
| C4 | C4 was lowered (P<0.001) in serum of term pre-eclamptics | [212] |
| C4d | Placental immunochemistry showed that C4d was rarely present in placentas from healthy controls (3%), whereas it was observed in 50% of placentas obtained from preeclamptic women (P=0.001) | [190] |
| C5a | The mean cord plasma C5a concentration was higher in patients with PET (8.3 ng/ml ± 1.71) than normal women (3.2 ng/ml ± 0.35) P < 0.01) | [192] |
| C5b-9 | Severe preeclampsia was associated with marked elevations in urinary C5b-9 (median and interquartile range, 4.3 (1.2–15.1) ng/mL) relative to subjects with chronic hypertension and healthy controls (P<0.0001). | [213] |
| C6 | Novel evidence that genetic variations in complement genes C6 and MASP1- were associated with preeclampsia risk | [198] |

The complement cascade may be activated in three main ways Fig 4), known as classical, alternative or lectin pathways [130; 186; 188; 199; 200]. Complement activation by the classical, alternative or lectin pathway results in the generation of split products C3a, C4a and C5a with pro-inflammatory properties.

**Figure 4.**
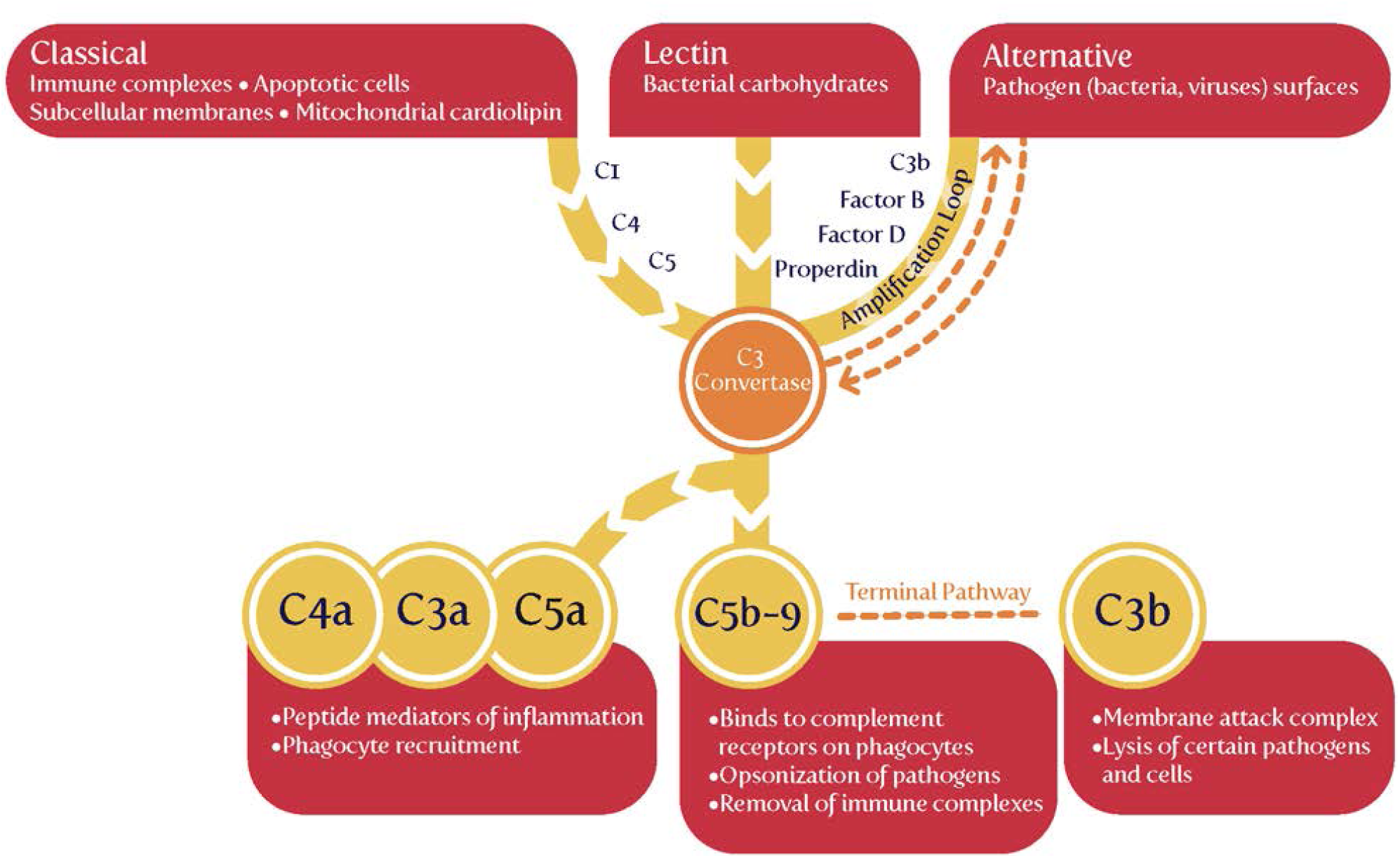
The complement system (based on figures in [135; 188]).

Because both innate and adaptive immunity can activate elements of the downstream complement system, it is hard to be definitive, but there is some evidence that elements such as Ba and Bb (the latter of known structure [201]) are selectively released during infection, very much upstream and in the alternative pathway [188; 199; 200; 202-204]. Most importantly (Table 1), while probably not a specific serum marker, there is considerable evidence that Bb levels <u>are</u> increased in PE, arguably providing further evidence for a role of infectious agents in the aetiology of PE.

We might also note that C1q^-/-^ mice shows features of PE [214], consistent with the view that lowering levels of anti-inection response elements of the complement system leads to PE, consistent again with an infectious component to PE.

## Induction of tolerance by exposure to antigens and our main hypothesis: roles of semen and seminal plasma

A number of groups (e.g. [100; 128; 215-218]) have argued for a crucial role of semen in inducing maternal immunological protection, and this is an important part of our own hypothesis here. The second component, however, is a corollary of it. **If it is accepted that semen can have beneficial effects, it may also be that <u>in certain cases it can also have harmful effects.</u>** Specifically, we rehearse the fact that semen is not sterile, and that it can be a crucial source of the microbes that may, over time, be responsible for the development of PE (and indeed other disorders of pregnancy, some of which we rehearse).

Semen consists essentially of the sperm cells suspended in a fluid known as seminal plasma [219]. Seminal plasma contains many components [220; 221], such as transforming growth factor β (TGF-β) [216; 222-224], and there is much evidence that a number of them are both protective and responsible for inducing the immune tolerance observed in pregnancy. Thus, in a key paper on the issue, Robertson and colleagues state, “TGFβ has potent immune-deviating effects and is likely to be the key agent in skewing the immune response against a Type-1 bias. Prior exposure to semen in the context of TGFβ can be shown to be associated with enhanced fetal/placental development late in gestation. In this paper, we review the experimental basis for these claims and propose the hypothesis that, in women, the partner-specific protective effect of insemination in pre-eclampsia might be explained by induction of immunological hyporesponsiveness conferring tolerance to histocompatibility antigens present in the ejaculate and shared by the conceptus” [128].

TGFβ and prostaglandin E (also prevalent in seminal fluid [225]) are potent Treg cell-inducing agents, and coitus is one key factor involved in expanding the pool of inducible Treg cells that react with paternal alloantigens shared by conceptus tissues [226-229].

Both in humans and in agricultural practice, semen may be stored with our without the seminal fluid (in the latter cases, the sperm have been removed from it and they alone are used in the insemination). However, a number of papers have shown very clearly that it is the seminal fluid itself that contains many protective factors, not least in improving the likelihood of avoiding adverse pregnancy outcomes [128; 177; 230; 231]. Thus semen is the preferred substrate for inducing immunotolerance and hence protection against PE.

## Evidence from epidemiology – semen can be protective against PE

As well as those (such as pre-existing diseases such as hypertension and diabetes [232; 233], that we covered previously [32]), there are several large-scale risk (or anti-risk) factors that correlate with the incidence of pre-eclampsia. They are consistent with the idea that a woman’s immune system adapts slowly to (semen) proteins from a specific male partner [128; 215; 216], and that the content of semen thus has major phenotypic effects well beyond its donation of (epi)genetic material. We believe that our hypothesis about the importance of semen in PE has the merit of being able to explain each of them in a simple and natural way:

1. The first pregnancy with any given partner means an increased susceptibility to PE [5; 234; 235]
2. Conception early in a new relationship means an increased susceptibility to PE [236-238]
3. Conception after using barrier contraceptives means an increased susceptibility to PE [237; 239; 240]
4. Conception after using non-barrier methods or after a long period of cohabitation means a decreased susceptibility to PET [215; 237]
5. Donor egg pregnancies have a hugely inflated chance of PET [235; 241-243]
6. Pre-eclampsia in a first pregnancy increases its likelihood in subsequent pregnancies [244]
7. Oral sex with the father is protective against PE in a subsequent pregnancy [245; 246]
8. Age is a risk factor for PE [247-251].
9. Donor sperm pregnancies (artificial insemination) are much more likely to lead to PE [246; 252-255]

We consider each in turn (Figure 5).

**Figure 5.**
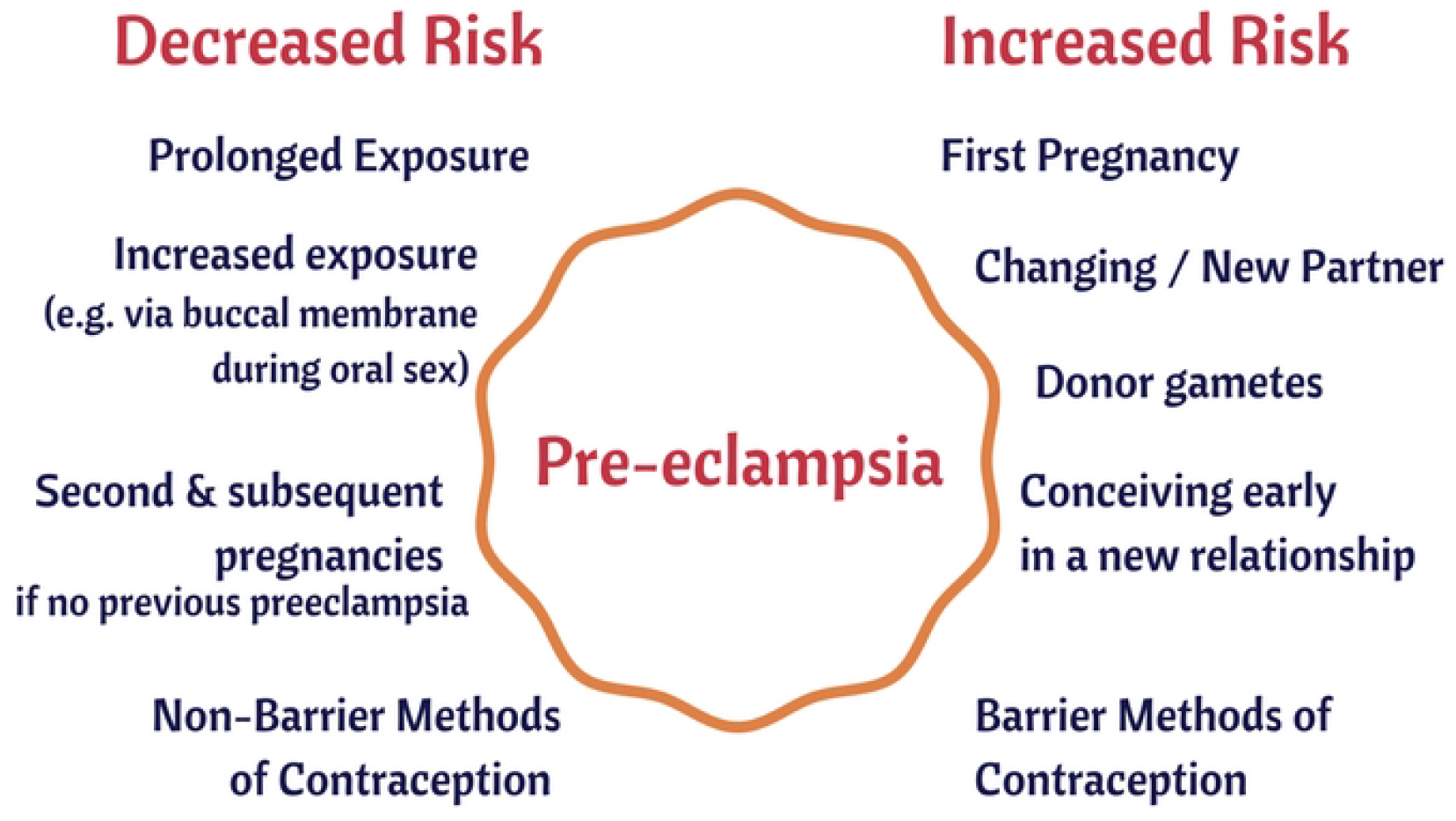
Some epidemiological risk factors for pre-eclampsia.

### The first pregnancy with any given partner means an increased susceptibility to PE

This is extremely well established (e.g. [5; 54; 149; 232; 234; 235; 256-263]). Thus, Duckitt and Harrington [232] showed nulliparity to have a risk ratio (over pregnant women with previous pregnancies) of 2.91 (95% CI 1.28-6.61). Luo *et al*. [259] find an odds ratio of 2.42 (95% CI 2.16-2.71) for PE in primiparous vs multiparous women, while Deis and colleagues found the OR to be 2.06 (CI 1.63 – 2.60), p = 0.0021. Dildy and colleagues [264] summarise several studies, including a very large one by Conde-Agudelo and Belizán [265] (RR 2·38; 95% CI 2·28-2·49), while the meta-analysis of English and colleagues [262] gives a risk ratio for nulliparity of 2.91 (CI 1.28 – 6.61). The consistency of each of these studies allows one to state with considerable confidence that there is a 2 – 3-fold greater chance of PE with a first baby.

However, an additional and key clue here is not simply (and maybe even not mainly) that it is just being nulliparous (i.e. one’s first pregnancy) but that it is primipaternity – one’s first pregnancy <u>with a given father</u> – that leads to an increased susceptibility to PE [183; 266-277] (cf. [278]). Changing partners effectively ‘resets the clock’ such that the risk with a new father is essentially as for first pregnancies. Thus, Lie *et al*. [279] noted that if a woman becomes pregnant by a man who has already fathered a pre-eclamptic pregnancy in a different woman her increased risk of developing pre-eclampsia is 1.8-fold (CI 1.2-2.6). This is far greater than the typical incidence of PE, even for nulliparous women. The equivalent figure in the study of Lynch and colleagues [183] was RR = 5.1, 95% CI, 1.6 to 15. The strong implication of all of this is that the father can have bad effects but that some kind of ‘familiarity’ with the partner is protective [275], the obvious version – and that more or less universally accepted – being an <u>immunological</u> familiarity (i.e. tolerance). Note, however, that this is when the pregnancy goes to term: a prior birth confers a strong protective effect against preeclampsia, whereas a prior abortion confers only a weaker protective effect [235].

### Conception early in a new relationship means an increased susceptibility to PE

The idea that conception early in a new relationship means an increased susceptibility to PE follows immediately from the above. The landmark studies here are those of Robillard and colleagues [236], of Einarsson and colleagues [237], and of Saftlas and colleagues [238],

Robillard *et al*. [236] studied 1011 consecutive mothers in an obstetrics unit. The incidence of pregnancy-induced hypertension (PIH) was 11.9% among primigravidae, 4.7% among same-paternity multigravidae, and 24.0% among new-paternity multigravidae. For both primigravidae and multigravidae, the length of (sexual) cohabitation before conception was inversely related to the incidence of PIH (P < 0.0001).

Einarsson and colleagues [237] studied both the use of barrier methods and the extent of cohabitation prior to pregnancy. For those (allegedly…) using barrier methods before insemination, the odds radio for PE when prior cohabitation was only 0-4 months versus the odds ratio for PE: normotensive was 17.1 (CI 2.9-150.6), versus 1.2 (CI 0.1-11.5) when the period of cohabitation was 8-12 months, and 1.0 for periods of cohabitation exceeding one year.

Saftlas *et al*. [238] recognised that parous women who change partners before a subsequent pregnancy appear to lose the protective effect of a prior birth. In a large study (mainly based around calcium supplementation), they noted that women with a history of abortion who conceived again with the same partner had nearly half the risk of preeclampsia (adjusted odds ratio = 0.54, 95 percent confidence interval: 0.31, 0.97). In contrast, women with an abortion history who conceived with a new partner had the same risk of preeclampsia as women without a history of abortion (adjusted odds ratio = 1.03, 95 percent confidence interval: 0.72, 1.47). Thus, the protective effect of a prior abortion operated only among women who conceived again with the same partner

### Conception after using barrier contraceptives means an increased susceptibility to PE

A prediction that follows immediately from the idea that paternal antigens in semen (or seminal fluid) are protective is that the regular use of barrier methods will lower maternal exposure to them, and hence increase the likelihood of PE. This too is borne out [237; 239; 240]. Thus Klonoff-Cohen and colleagues found a 2.37-fold (CI 1.01-5.58) increased risk of preeclampsia for users of contraceptives that prevent exposure to sperm. A dose-response gradient was observed, with increasing risk of preeclampsia for those with fewer episodes of sperm exposure. Similarly, Hernández-Valencia and colleagues [240] found that the odds ratio for preeclampsia indicated a 2.52-fold (CI 1.17-5.44, P < 0.05), increased risk of preeclampsia for users of barrier contraceptives compared with women using nonbarrier contraceptive methods.

### Conception after using non-barrier methods or after a long period of cohabitation means a decreased susceptibility to PE

This is the flip side of the studies given above (e.g. [236-238]). It is clear that maternal–fetal HLA sharing is associated with the risk of preeclampsia, and the benefits of long-term exposure to the father’s semen, while complex [280], seem to be cumulative [281]. Thus, short duration of sexual relationship was more common in women with preeclampsia compared with uncomplicated pregnancies (≤ 6 months 14.5% versus 6.9%, adjusted odds ratio (aOR) 1.88, 95% CI 1.05–3.36; ≤3 months 6.9% versus 2.5%, aOR 2.32, 95% CI 1.03–5.25 [282]. Oral contraceptives are somewhat confounding here, in that they may either be protective or a risk factor depending on the duration of their use and the mother’s physiological reaction to them [283].

### Donor egg pregnancies have a hugely inflated chance of PE

If an immunological component is important to PE (as it evidently is), it is to be predicted that donor egg pregnancies are likely to be at much great risk of PE, and they are (e.g. [235; 241-243; 284-288]) (and also of pre-term birth [289]). Thus, Letur and colleagues [241; 242] found that preeclampsia was some fourfold more prevalent using donated eggs (11.2% vs. 2.8%, P < 0.001), while Tandberg and colleagues [235] found that various ‘assisted reproductive technologies’ had risk ratios of 1.3 (1.1–1.6) and 1.8 (1.2–2.8) in second and third pregnancies, respectively. Pecks and colleagues studied pregnancy-induced hypertension (PIH, not just PE) and found that the calculated odds ratio for PIH after oocyte donation, compared to conventional reproductive therapy, was 2.57 (CI 1.91–3.47), while the calculated odds ratio for PIH after oocyte donation, compared to other women in the control group, was 6.60 (CI 4.55–9.57). Stoop and colleagues [290] found a Risk Ratio of 1.502 (CI 1.024-2.204) for PIH. In a study by Levron and colleagues [291], adjustment for maternal age, gravidity, parity, and chronic hypertension revealed that oocyte donation was independently associated with a higher rate of hypertensive diseases of pregnancy (P < 0.01). In a twins study, Fox and colleagues [292] found, on adjusted analysis, that the egg donation independently associated with preeclampsia (aOR 2.409, CI 1.051-5.524). The meta-anaysis of Thomopoulos and colleagues [293] gave a risk ratio for egg donation of 3.60 (CI 2.56–5.05) over controls, a value similar to that of Blázquez and colleagues [294]. Finally, a recent meta-analysis by Masoudian and colleagues [287] found that that the risk of preeclampsia is considerably higher in oocyte-donation pregnancies compared to other methods of assisted reproductive technology (odds ratio, 2.54; CI 1.98-3.24; P < 0.0001) or to natural conception (odds ratio, 4.34; CI 3.10-6.06; P < 0.0001). The incidence of gestational hypertension and preeclampsia was significantly higher in ovum donor recipients compared with women undergoing autologous IVF (24.7% compared with 7.4%, P < 0.01, and 16.9% compared with 4.9%, P < 0.02 [295]. All of these are entirely consistent with an immune component being a significant contributor to PE. One obvious question pertains to whether the use of antibiotics assists the progression of IVF. Unfortunately this question has been little researched in humans [296].

### PE in a first pregnancy increases its likelihood in subsequent pregnancies

This too is well established: a woman who has had preeclampsia has an increased risk of preeclampsia in subsequent pregnancies [263; 297], especially if suffering from hypertension [298]. This may be seen as relatively unsurprising, and of course bears many explanations, and the increased risks can be very substantial [244]. In the overall analysis of English and colleagues [262], the risk ratio was 7.19 (CI 5.85–8.83). Other examples give the recurrence risk, overall, as some 15% to 18% [263]. The risk of recurrent preeclampsia is inversely related to gestational age at the first delivery, and in the study of Mostello and colleagues [299] was 38.6% for 28 weeks’ gestation or earlier, 29.1% for 29-32 weeks, 21.9% for 33-36 weeks, and 12.9% for 37 weeks or more. Low birthweight in the first pregnancy is an independent predictor of PE in the second: birth weight below the tenth percentile in the first delivery accounted for 10% of the total cases of preeclampsia in the second pregnancy and 30% of recurrent cases [300]. From the perspective developed here, the suggestion is that whatever is responsible for PE in one pregnancy can ‘live on’ in the mother and afflict subsequent ones. One thing that can ‘live on’ is a dormant microbial community.

### Oral sex with the father is protective against pre-eclampsia in a subsequent pregnancy

Oral sex (with the father of one’s baby) protects against pre-eclampsia [245; 246] (p = 0.0003), arguably because exposure to the paternal antigens in the seminal fluid have a greater exposure to the blood stream via the buccal mucosa than they would via the vagina. This is a particularly interesting (and probably unexpected) finding, that is relatively easily understood from an immunological point of view, and it is hard to conceive of alternative explanations. (Note, however, that in the index study [245], the correlation or otherwise of oral and vaginal sex was not reported, so it is not entirely excluded that more oral sex also meant more vaginal sex.)

### Age is a risk factor for PE

Age is a well known risk factor for PE [247-251], and of course age is a risk factor for many other diseases, so we do not regard this as particularly strong evidence for our ideas. However, we have included it in order to note that age-associated microbial dysbiosis promotes intestinal permeability, systemic inflammation, and macrophage dysfunction [301].

### Donor sperm pregnancies (artificial insemination) are much more likely to lead to PE

Finally, here, turning again to the father, it has been recognised that certain fathers can simply be ‘dangerous’ in terms of their ability to induce PE in those who they inseminate [277; 302]. By contrast, if immunotolerance to a father builds up slowly as a result of cohabitation and unprotected sex, a crucial prediction is that donor sperm pregnancies will not have this property, and should lead to a much greater incidence of PE. This is precisely what is observed [246; 252-255; 284].

In an early study [252], Need and colleagues observed that the overall incidence of PE was high (9.3%) in pregnancies involving artificial insemination by donor (AID) compared with the expected incidence of 0.5-5.0%. The expected protective effect of a previous pregnancy was not seen, with a 47-fold increase in PE (observed versus expected) in AID pregnancies after a previous full-term pregnancy. That is a truly massive risk ratio.

Smith and colleagues [253] compared the frequency of PE when AI was via washed sperm from a partner or a donor, finding a relative risk for PE of 1.85 (95% CI 1.20-2.85) for the latter, and implying that the relevant factor was attached to (in or on) the sperm themselves.

In a similar kind of study, Hoy and colleagues found [254], after adjusting for maternal age, multiple birth, parity and presentation, that ‘donor sperm’ pregnancies were more likely to develop preeclampsia (OR 1.4, 95% CI 1.2–1.8).

Salha and colleagues [284] found that the incidence of pre-eclampsia in pregnancies resulting from donated spermatozoa was 18.2% (6/33) compared with 0% in the age-and parity-matched partner insemination group (P < 0.05).

Wang *et al*. [303] found that the risk of pre-eclampsia tripled in those never exposed to their partner’s sperm, i.e, those treated with intracytoplasmatic sperm injection done with surgically obtained sperm.

In a study of older women, Le Ray and colleagues [304] noted that the pre-eclampsia rate differed significantly between various groups using assisted reproductive technology (3.8% after no IVF, 10.0% after IVF only and 19.2% after IVF with oocyte donation, P < 0.001).

Davis and Gallup reviewed what was known in 2006 [255], particularly from an evolutionary point of view, concluding that one interpretation of PE was that it was the mother’s way of removing ‘unsuitable’ fetuses. This does not sit easily with the considerable mortality and morbidity associated with PE pre-delivery, especially in the absence of treatment. However, Davis and Gallup [255] did recognise that “pregnancies and children that result from unfamiliar semen have a lower probability of receiving sufficient paternal investment than do pregnancies and children that result from familiar semen”, and that is fully consistent with our general thinking here. Bonney draws a similar view [162], based on the ‘danger’ model [156; 158], that takes a different view from that of the ‘allograft’ or ‘self-nonself discrimination’ model. In the ‘danger model’, the decision to initiate an immune response is based not on discrimination between self and non-self, but instead is based on the recognition of ‘danger’ (abnormal cell death, injury or stress). One such recognition is the well-established recognition of microbes as something likely to be causative of undesirable outcomes.

In the study of González-Comadran and colleagues, [305], conception using donor sperm was again associated with an increased risk of preeclampsia (OR 1.63, 95% CI 1.36–1.95).

Thomopoulos and colleagues carried out two detailed and systematic reviews [293; 306]; the latter [293] covered 7,038,029 pregnancies (203,375 following any invasive ART) and determined that the risk of PE was increased by 75% (95% CI, 50%–103%).

Overall, these studies highlight very strongly indeed that the use of unfamiliar male sperm is highly conducive to PE relative to that of partner’s sperm, especially when exposure is over a long period. We next turn to the question of why, in spite of this, we also see PE even in partner-inseminated semen, as well as more generally.

## Evidence from epidemiology–semen can be harmful and can contribute strongly to PE

In our previous review [32], we rehearsed the evidence for a considerable placental and vaginal microbiome, but did not discuss the semen microbiome at all. To repeat, therefore, the particular, and essentially novel, part of our hypothesis here is that **if it is accepted that semen (and seminal plasma) can have <u>beneficial</u> effects, it should also be recognised that in certain cases it can also have <u>harmful</u> effects.** In particular, we shall be focussing on its microbial content (we ignore any epigenetic effects [307]). We note that this idea would fit easily with the recognition that as well as inducing tolerance to paternal antigens, exposures to the father’s semen can build tolerance (immunity) to its microbes, thereby decreasing the risk of PE. However, microbes and their associated PAMPs are well known to be highly inflammatory, whether or not they are reproducing, and we consider that it is this that is likely the particular driver of the sequelae observable in PE.

## Microbes associated with pre-eclampsia

The female’s urogenital microbiome is important in a number of pregnancy disorders [44; 308-310]. Specifically, we previously found many examples in which microbes are associated with PE, and we here update the CC-BY-licensed Table 2 thereof [32] as Table 2 here:

**Table 2.**
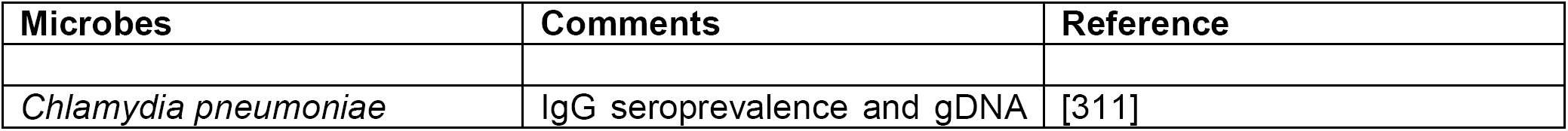
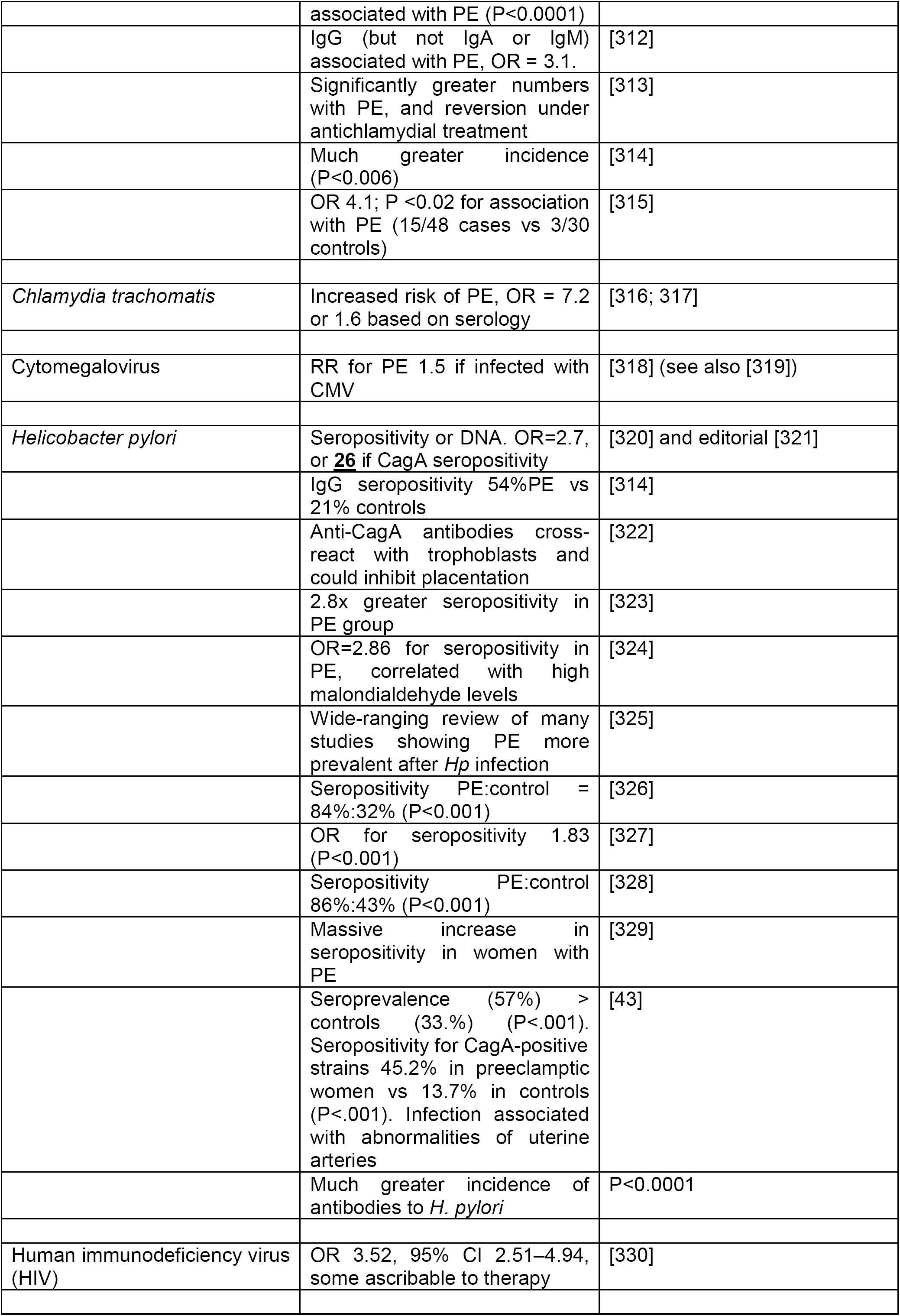
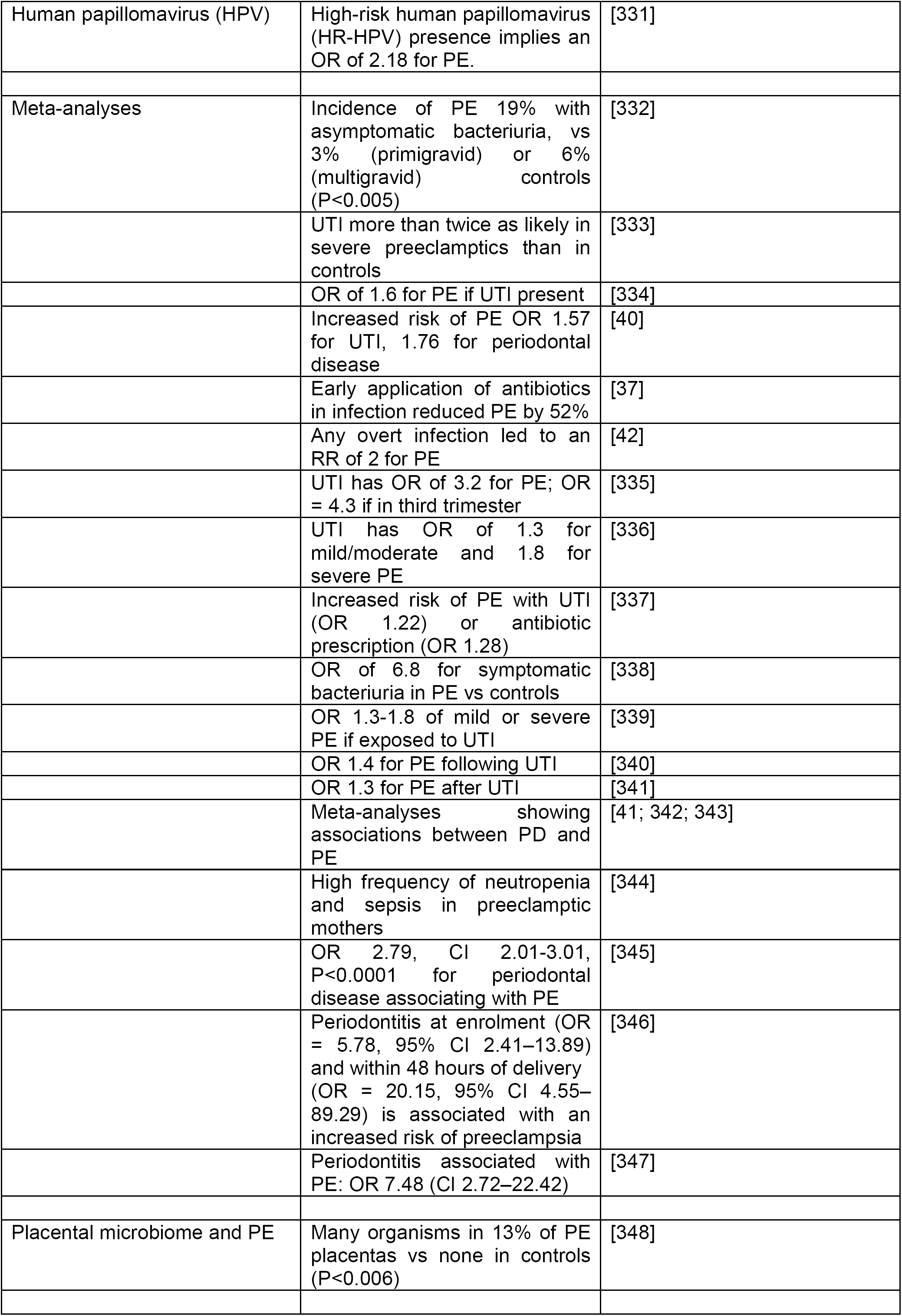
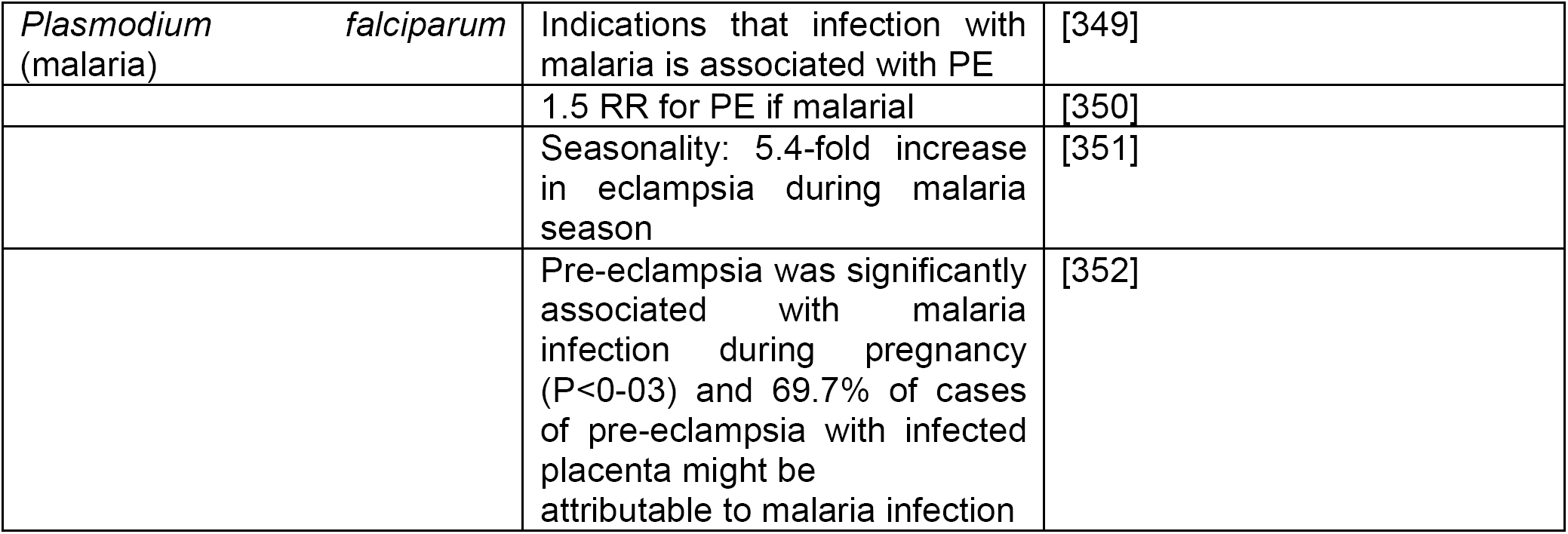
Many studies have identified a much greater prevalence of infectious agents in the blood or urine or gums of those exhibiting PE than in matched controls

## Microbiology of semen

Semen itself is very far from being sterile, even in normal individuals, with infection usually being defined as 10^3^ organisms.mL^−1^ semen [353]. Of course the mere existence of sexually transmitted diseases implies strongly that there is a seminal fluid (or semen) microbiome that can vary substantially between individuals, and that can contribute to infection (e.g. [354-356]), fertility [354] (and see below), and any other aspect of pregnancy [357], or even health in later life [358].

It is logical to start here with the observation that semen <u>is</u> a source of microbes from the fact that there are a great many sexually transmitted infectious diseases for which it is the vehicle. Table 3 summarises some of these.

**Table 3.** Organisms of well-known sexually transmitted diseases that have been associated with semen

| Organism (disease) | Comments | References |
| --- | --- | --- |
| *Chlamydia trachomatis* | Effects on fertility | [359] |
|  | 32% prevalence in infertile couples | [360] |
| Human Immunodeficiency Virus (AIDS) | Many examples of seminal transmission via unprotected sex | [361-366] |
| *Neisseria gonorrhoeae* (gonorrhoea) | Gonorrhea actually means 'flow of semen' | [367] |
|  | Survives being frozen in semen used for artificial insemination | [368] |
|  | Many anti-gonococcal antibodies also present | [369] |
|  | Same strains in urine and semen; likely origin in urethra | [370] |
| *Treponema pallidum* (syphilis) | Infectivity of semen | [371] |
|  | More than half (twelve out of twenty) of the women classified | [372] |
|  | as proved and probably syphilitic had mild to moderate PE |  |
|  | Co-infection of syphilis and HIV in men having sex with men | [373] |
|  | Both syphilis and preeclampsia contribute to stillbirths in sub-Saharan Africa | [374] |

Notwithstanding the difficulties of measurement [375], there is, in particular, a considerable literature on fertility [376], since infertile males tend to donate sperm for assay in fertility clinics, and infection is a common cause of infertility (e.g. [353] and Table 4). Note that ‘infertility’ is not always an absolute term: pregnancies result in 27% of cases of treated ‘infertile’ couples followed up after trying to conceive for 2y, and with oligozoospermia as the primary cause of infertility [377]. Most studies involve bacteria (bacteriospermia). Papers on this and other microbial properties of semen beyond STDs include those in Table 4.

**Table 4.** Some examples of the semen microbiome and reproductive biology

| Study | Organisms | References |
| --- | --- | --- |
| Complementarity between partners | Many. *Gardnerella vaginalis* in female partners was significantly related to inflammation in male genital tracts | [378] |
| Fertility | Many microbiological changes as a function of fertility (more microbes correlate with lower fertility) | [353; 357; 377; 379-423] |
| General microbiology | 552 different microbes in 182 samples out of 201 tested, simply plating 10 μL of semen | [424] |
|  | Microbes in 36/37 samples | [425] |
|  | Review | [426] |
|  | 35% of samples had microbes | [427] |
| IVF | No positive antibiotic effect | [428] |
| LPS and protection by probiotic lactobacilli | (purified LPS) | [429] |
| Review | Many microbes | [415; 430; 431] |
| Semen quality | *Ralstonia* increased in low-quality sperm | [432] |
| Viral infection | Ebola virus | [433-436] |
|  |  | [437] |
|  | HIV | [438-440] |
|  | Zika virus |  |

We deliberately avoid discussing mechanisms in any real detail here, since our purpose is merely to show that semen <u>is</u> commonly infected with microbes, whose presence might well lead to preeclampsia. However, we were very struck by the ability of *E. coli* and other organisms [410; 420; 441] actually to immobilise sperm (e.g. [442-445]). As with amyloidogenic blood clotting [446; 447], bacterial LPS [136] may be a chief culprit [429]. The Gram-positive equivalent, lipoteichoic acid (LTA), is just as potent in the fibrinogen-clotting amyloidogen assay [448], but while Gram-positives can also immobilise sperm [449; 450], the influence of purified LTA on sperm seems not to have been tested.

Another prediction from this analysis is that since infection is a significant cause of <u>both</u> infertility and PE (and it may account for 15% of infertile cases [353; 443]), we might expect to see some correlations between them. Although one might argue that anything seen as imperfect ‘background’ health or subfecundity might impinge on the incidence of PE, the risk ratio for PE in couples whose infertility had an unknown basis was 5.61 (CI 3.3-9.3) in one study in Aberdeen [451] and 1.29 (CI 1.05–1.60) in another in Norway [452]. Time to pregnancy in couples may be used (in part) as a surrogate for (in)fertility and is associated with a variety of poor pregnancy outcomes [453]; in this case, the risk ratio for PE for TTP exceeding 6 months was 2.47 (CI 1.3-4.69) [454]. Given the prevalence of infection in infertile sperm (Table 4), and the frequency of infertility (10% in the Danish study [453], which defined it as couples taking a year or more to conceive), it seems reasonable to suggest that microbiological testing of semen should be done on a more routine basis. It would also help to light up any relationships between the microbiological properties of sperm and the potentially causal consequence of increased PE risk.

More quantitatively, and importantly intellectually, if infection is seen as a major cause of PE, as we argue here, and it is known that infection is a cause of infertility, then one should suppose that infertility, and infertility caused by infection, should be at least as common, and probably more common than is PE, and this is the case, adding some considerable weight to the argument. Indeed, if PE was much more common than infertility or even infection, it would be much harder to argue that the latter was a major cause of the former. In European countries ~10–15% of couples are afflicted by infertility [353; 453], and in some 60% of cases infection or a male factor is implicated [353]. In some countries, the frequency of male infertility is 13-15% http://bionumbers.hms.harvard.edu/bionumber.aspx?id=113483&ver=0 or higher [455], and the percentage of females with impaired fecundity has been given as 12.3% https://www.cdc.gov/nchs/fastats/infertility.htm. These kinds of numbers would imply that 6-9% of couples experience infection-or male-based infertility, and this exceeds the 3-5% incidence of PE.

In a similar vein, antibiotics, provided they can get through the relevant membranes [456-458], should also have benefits on sperm parameters or fertility if a lack of it is caused by infection, and this has indeed been observed (e.g. [407; 423; 459]).

## Roles of the prostate and testes

In the previous review, we focussed on the gut, periodontitis, and the urinary tract of the mother as the main source of organisms that might lead to PE. Here we focus on the male, specifically the prostate and the testes, given the evidence for how common infection is in semen. The main function of the prostate gland is to secrete prostate fluid, one of the components of semen. Thus, although it is unlikely that measurement have regularly been done to assess any relationship between this and any adverse effects of pregnancy, it was of interest to establish whether it too is likely to harbour microbes. Indeed, such ‘male accessory gland infection’ is common [460-464]. In some cases, the origin is probably periodontal [465]. Recent studies have implicated microbial pattern-recognition receptors, especially Toll-like receptors (TLRs), as well as inflammatory cytokines and their signalling pathways, in testicular function, implying an important link between infection/inflammation and testicular dysfunction [466]. The testes are a common and important site of infection in the male [467; 468], and even bacterial LPS can cause testitis [469]. Similarly, infection (especially urinary tract infection) is a common cause of prostatitis [470-480]. Finally, prostatitis is also a major cause of infertility [460; 461; 463]. Such data contribute strongly to the recognition that semen is not normally going to be sterile, consistent with the view that it is likely to be a major originating cause of the infections characteristic of PE.

## Microbial infections in spontaneous abortions, miscarriages and pre-term birth

Our logic would also imply a role for (potentially male-derived) microbes in miscarriages and spontaneous abortions. A microbial component to these seems well established for both miscarriages [481-483] and spontaneous abortions [484-489]. Of course the ability of *Brucella abortus* to induce abortions in domesticated livestock, especially cattle (and occasionally in humans), is well known [490-492]; indeed, bacteriospermia is inimical to fertilisation success [493], and nowadays it is well controlled in livestock by the use of vaccines [494] or antimicrobials [493]. Indeed, stored semen is so widely used for the artificial insemination of livestock in modern agriculture that the recognition that semen is not sterile has led to the routine use of antibiotics in semen ‘extenders’ (e.g. [495-498]).

The same general logic is true for infection as a common precursor to pre-term birth (PTB) in the <u>absence</u> of PE, where it is much better established (e.g. [499-533]). It arguably has the same basic origins in semen.

Although recurrent pregnancy loss is usually treated separately from infertility (where the role of infection is reasonably well established) it is possible that in many cases it is, like PE, partly just a worsened form of an immune reaction, with both sharing similar causes (including the microbial infection of semen). There is in fact considerable evidence for this (e.g. [120; 413; 534-548]). Of course it is not unreasonable that poor sperm quality, that may be just sufficient to <u>initiate</u> a pregnancy, may ultimately contribute to its premature termination or other disorders of pregnancy, so this association might really be expected. It does, however, add considerable weight to the view that a more common screening of the male than presently done might be of value [549] in assessing a range of pregnancy disorders besides PE. In particular, it seems that infection affects motility (see above), and that this in turn is well correlated [541] with sperm DNA fragmentation and ultimate loss of reproductive quality.

Amyloids in semen are known to enhance HIV infectivity [550]. According to our own recent experimental analyses, they may be caused by bacterial lipopolysaccharide (LPS) [446; 447] or lipoteichoic acid [448]. We note too that the sperm metabolome also influences offspring, e.g. from obese parents [551], and that many other variables are related to sperm quality, including oxidative stress [552-559]. Thus it is entirely reasonable to see semen as a cause of problems as well as benefits to an ensuing pregnancy.

## Microbial effects on immunotolerance

If our thesis is sound, one may expect to find evidence for the effects of microbes on the loss of immunotolerance in <u>other</u> settings. One such is tolerance to dietary antigens, of which gluten, a cause of coeliac disease, is pre-eminent. Recently, evidence has come forward that shows a substantial effect of a reovirus in lowering the immunotolerance to gluten in a mouse model of coeliac disease, and thereby causing inflammation [560; 561]. Interestingly, pregnancies in women with coeliac disease were considerably more susceptible to pre-term birth and other complications than were controls [562-569], especially when mothers were not on a gluten-free diet. Similarly, pre-eclamptic pregnancies led to a much (4-fold) higher likelihood of allergic sensitisation in the offspring [570] The roles of hygiene, the microbiome and disease are a matter of considerable current interest (e.g. [571]).

It was consequently logical to see if intolerance to peanut antigen was also predictive of PE, but we could find no evidence for this. Again, however, in a study [572] in which PE had roughly its normal prevalence, mothers experiencing it were significantly more likely to give birth to children with increased risk of asthma, eczema, and aeroallergen and food allergy.

## Effects of vaccination on pregnancy outcomes, including pre-eclampsia

We noted above (and again below) that the evidence for a role of microbes in pre-term birth (PTB) is overwhelming (also reviewed in [32]). From an immunological point of view, there seems to be a hugely beneficial outcome of vaccination against influenza in terms of lowering pre-term birth [573-578] (cf. [579]) or stillbirth [580]. (PE was not studied, save in [581] where the risk ratio of vaccination (0.484, CI 0.18–1.34) implied a marginal benefit. There do not seem to be any safety issues, either for influenza vaccine [580-601] or for other vaccines [593] such as those against pertussis [602-604] or HPV [605].

As well as miscarriage and pre-term birth, other adverse pregnancy outcomes studied in relation to vaccine exposure [606] include intrauterine growth restriction (IUGR). IUGR frequently presents as the fetal phenotype of pre-eclampsia, sharing a common aetiology in terms of poor placentaton in early pregnancy [607]. These other adverse events have been scored more frequently than has been PE, and Table 5 summarises the evidence for a protective effect of vaccines, though it is recognised that there is the potential for considerable confounding effects (e.g. [600; 608]).

**Table 5.** Protective events of vaccines against various adverse pregnancy outcomes

| Adverse event | Risk or Odds ratio (95% Confidence interval) of vaccinated: unvaccinated | Reference |
| --- | --- | --- |
| *Pre-term birth* | [OR = 0.39 (0.18-0.83)] | [574] |
|  | 0.56 (0.45-0.70) | [576] |
|  | 0.60 0.38–0.94<br>0.28 (0.11–0.74) during epidemic | [573] |
|  | 0.63 (0.47–0.84) | [575] |
| IUGR | 0.15 (0.02-0.94) | [574] |
|  | 0.36 (0.17–0.78) | [592] |
|  | 0.31 (0.13–0.75) | [573] |
|  | 0.63 (0.4–1.0) | [609] |
| Stillbirth | 0.73 (0.55-0.96) | [580] |

There are no apparent benefits of vaccine-based immunisation vs recurrent miscarriage [610; 611].

Unrelated to the present question, but very interesting, is the fact that the risk of RA <u>for men</u> was higher among men who fathered their first child at a young age (p for trend < 0.001) [612]. This is consistent with the fact that its prevalence in females is 3.5 times higher, and that it has a microbial origin [613-616].

## General or specific?

The fact that vaccination against organisms not usually associated with adverse pregnancy outcomes is protective can be interpreted in one (or both) of two ways, i.e. that the vaccine is unselective in terms of inhibiting the effects of its target organism, or the generally raised level of <some kind of> immune response is itself protective. Data to discriminate these are not yet to hand.

In a similar vein, the survival of the host in any ‘battle’ between host and parasite (e.g. microbe) can be effected in one or both of two main ways: (i) the host invokes antimicrobial processes such as the immune systems described above, or produces antimicrobial compounds, or (ii) the host modifies itself in ways that allow it to become tolerant to the presence of a certain standing crop of microbes. We consider each in turn.

## Antimicrobial components of human semen, a part of resistance in the semen microbiome

Antimicrobial peptides (AMPs) (http://aps.unmc.edu/AP/main.php [617]) are a well-known part of the defence systems of many animals (e.g. [618-627]) (and indeed plants [618; 628]), and are widely touted as potential anti-infectives (e.g. [629-631]). Their presence in the cells and tissues of the uterus, fetus and the neonate indicates an important role in immunity during pregnancy and in early life [625; 632-636]. Unsurprisingly, they have been proposed as agents for use in preventing the transmission of STDs [637; 638], and as antimicrobials for addition to stored semen for use in agriculture [639-643]. Our interest here, however, is around whether there are <u>natural</u> AMPs in human (or animal) semen, and the answer is in the affirmative. They include SLPI [627], SEVI [644], and in particular the semenogelins [645; 646]. HE2 is another antimicrobial peptide that resides in the epididymis [647; 648], while the human cathelicidin hCAP-18 in inactive in seminal plasma but is processed to the antimicrobial peptide LL-37 by the prostate-derived protease gastricsin [636; 649]. Thus it is clear that at least some of the reason that the semen microbiome is not completely unchecked is down to antimicrobial peptides. Stimulating their production, provided they are not also spermicidal, would seem like an excellent therapeutic option.

## Host tolerance to microbial pathogens

It is a commonplace that–for any number of systems biology reasons based on biochemical individuality [650]–even highly virulent diseases do not kill everyone who is exposed to them at the same level. As indicated above, this could be because the host is <u>resistant</u> and simply clears the infections; this is certainly the more traditional view. However, an additional or alternative contribution is because while host do not clear all of them they can develop <u>‘tolerance’</u> to them. This latter view is gaining considerable ground, not least since the work of Schneider, Ayres and colleagues [651] showing that a variety of *Drosophila* mutants with known genetic defects could differentially tolerate infection by *Listeria monocytogenes*. This concept of tolerance [652-659] is very important to our considerations here, since it means that we do indeed have well-established methods of putting up with microbes more generally, without killing them. It is consistent with clearly established evolutionary theory [660-662], and the relative importance of resistance and tolerance within a population affects host-microbe coevolution [663]. The concept of tolerance sits easily with the Matzinger model of danger/damage (e.g. [155; 157; 158; 160]), as well as the concept of a resident population of dormant microbes [33; 35; 36], and may indeed be seen in terms of a coevolution or mutualistic association [664; 665]. Some specific mechanisms are becoming established, e.g. the variation by microbes of their danger signal to promote host defence [666]. Others, such as the difference in the host metabolomes (that we reviewed [32]) as induced by resistance vs tolerance responses [658] may allow one to infer the relative importance of each. At all events, it is clear from the concept of dormancy that we do <u>not</u> kill all the intracellular microbes that our bodies harbour, and that <u>almost by definition</u> we must then tolerate them. As well as the established maternal immunotolerance of pregnancy, tolerance of microbes seems to be another hallmark of pregnancy.

## Sequelae of a role of infection in PE: microbes, molecules and processes

The chief line taken in our previous review [32] and herein is that this should be detectable by various means. Those three chief means involve detecting the microbes themselves, detecting molecules whose concentration changes as a result of the microbes (and their inflammatory components) being present, and detecting host processes whose activities have been changed by the presence of the microbes.

Previously [32], updated here (Table 2), we provided considerable evidence for the presence of microbes within the mother as part of PE. Here we have adduced the equally considerable evidence that in many cases semen is very far from being sterile, and that the source of the originating infection may actually be the father. Equally, we showed [32] that a long list of proteins that were raised (or less commonly lowered) in PE were equally changed by known infections, consistent with the view that PE also involved such infections, albeit at a lower level at which their overt presence could be kept in check. One protein we did not discuss was Placental Protein 13 or galectin 1, so we now discuss this briefly.

### Placental protein 13 (galectin 1)

Galectins are glycan-binding proteins that regulate innate and adaptive immune responses. Three of the five human cluster galectins are solely expressed in the placenta [667]. One of these, encoded by the *LGALS13* gene [668], is known as galectin-13 or Placental Protein 13 (PP13) [669]. Its β-sheet-rich ‘jelly-roll’ structure places it strongly as a galectin homologue [670]. It has a MW of ~16kDa (32kDa dimer [671]) and is expressed solely in the placenta [672] (and see http://www.proteinatlas.org/ENSG00000105198-LGALS13/tissue). A decreased placental expression of PP13 and its low concentrations in first trimester maternal sera are associated with elevated risk of preeclampsia [667; 673-675], plausibly reflecting poor placentation. By contrast, and consistent with the usual oxidative stress, there is increased trophoblastic shedding of PP13-immunopositive microvesicles in PE, starting in the second trimester, which leads to high maternal blood PP13 concentrations [667; 676]. Certain alleles such as promoter variant 98A-C predispose strongly to PE [677]. (Galectin-1 is also highly overexpressed in PE [678].) However, as with all the other proteomic biomarkers surveyed previously [32], galectins (including galectin-13 [679] http://amp.pharm.mssm.edu/Harmonizome/gene/LGALS13) are clear biomarkers of infection [680].

### Toll-like receptors (TLRs)

TLRs are among the best known receptors for ‘damage-associated molecular patterns’ such as LPS from Gram-negatives (TLR 4 [136; 681-683]), lipoteichoic acids (LTAs) from Gram-positives (TLR2 [684-695]) and viral DNA and its mimics (TLR3) [696].

As expected, they are intimately involved in disorders of pregnancy such as PE [165; 696-708]. Indeed the animal model for preeclampsia developed by Faas and colleagues [709] actually involves injecting an ultra-low dose of LPS into pregnant rat on day 14 of gestation. Overall, such data are fully consistent with the view that infection is a significant part of PE. In view of our suggestions surrounding the role of semen infection in PE it would be of interest to know if these markers were also raised in the semen of partners of women who later manifest PE. Sperm cells are well endowed with TLRs [466; 710-712]. However, we can find only one study showing that increased semen expression of TLRs is indeed observed during inflammation and oxidative stress such as occurs during infection and infertility [713]. A more wide-ranging assessment of TLR expression in sperm cells as a function of fertility seems warranted.

## Coagulopathies

Although we discussed this in the previous review [32], some further brief rehearsal is warranted, since coagulopathies are such a common feature of PE (references in [32]). Specifically, our finding that very low concentrations of cell wall products can induce amyloid formation during blood clotting [446; 448] has been further extended to recognise the ubiquity of the phenomenon in chronic, inflammatory diseases [447; 448; 616; 714-716]. Often, an extreme example gives strong pointers, and the syndrome with the highest likelihood of developing PE is antiphospholipid syndrome [717-721], which is also caused by infection [722-727] where the coagulopathies are also especially noteworthy [728-732]. Consequently, the recognition of PE as a amyloidogenic coagulopathy [32; 733-735] is significant.

## Antiphospholipid syndrome and cardiolipin

Antiphospholipid syndrome (APS) is an autoimmune disorder defined in particular by the presence high circulating titres of what are referred to as antiphospholipid antibodies (aPL) (e.g. [736]). Given that every human cell’s plasma membrane contains phospholipids, one might wonder how ‘antiphospholipid antibodies’ do not simply attack every cell. The answer, most interestingly, is that, despite the name, anticardiolipin antibodies, anti-β2–glycoprotein-I, and lupus anticoagulant are the main autoantibodies found in antiphospholipid syndrome [737].

In contrast to common phospholipids such as phosphatidylcholine, phosphatidylserine and phosphatidylethanolamine, cardiolipins (1,3-bis(sn-3’-phosphatidyl)-sn-glycerol derivatives) (see Figure 6 for some structures) are synthesised in [738-740] and essentially confined to mitochondria, and in particular the inner mitochondrial membrane, where they serve important functions in oxidative phosphorylation, apoptosis, and heart failure development [740-747]

**Figure 6.**
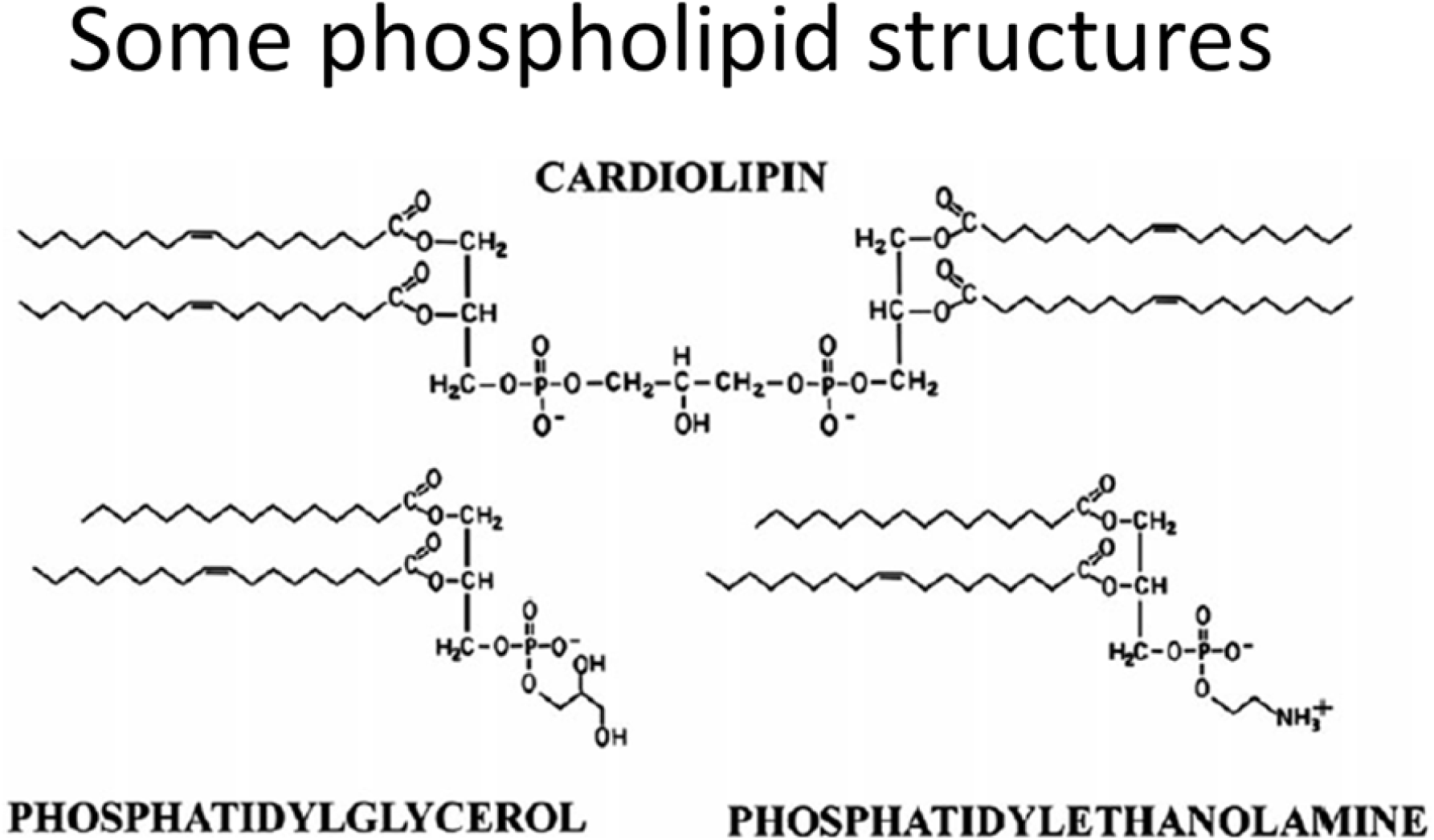
Some cardiolipin structures.

Overall, there seems to be little doubt that APS and aPL are the result of infection [722; 724-727; 748-750], and that, as with rheumatoid arthritis (see [613-616; 751]), the auto-immune responses are essentially due to molecular mimicry.

Now, of course, from an evolutionary point of view, mitochondria are considered to have evolved from (α-Proteo)bacteria [752-758] that were engulfed by a proto-eukaryote [759], and bacteria might consequently be expected to possess cardiolipin. This is very much the case for both Gram-negative and Gram-positive strains [760-764], with Gram-positive organisms typically having the greater content. Particularly significant, from our point of view, is that the relative content of cardiolipin among phospholipids increases enormously as (at least Gram-positive) bacterial cells become dormant [765].

Thus, the cardiolipin can come from two main sources: (i) host cell death that liberates mitochondrial products or (ii) invading bacteria (especially those that lay dormant and awaken). Serum ferritin is a cell death marker [766], and some evidence for the former source [767] (and see [768]) is that hyperferritinemia was present in 9% vs. 0% of APS patients and controls, respectively (p < 0.001), and that hyperferritinemia was present in 71% of catastrophic APS (cAPS) patients, and ferritin levels among this subgroup were significantly higher compared with APS–non-cAPS patients (816-847 ng/ml vs. 120-230 ng/ml, p < 0.001). One easy hypothesis is that both are due to invading bacteria, but cAPS patients also exhibit comparatively large amounts of host cell death (Figure 7).

**Figure 7.**
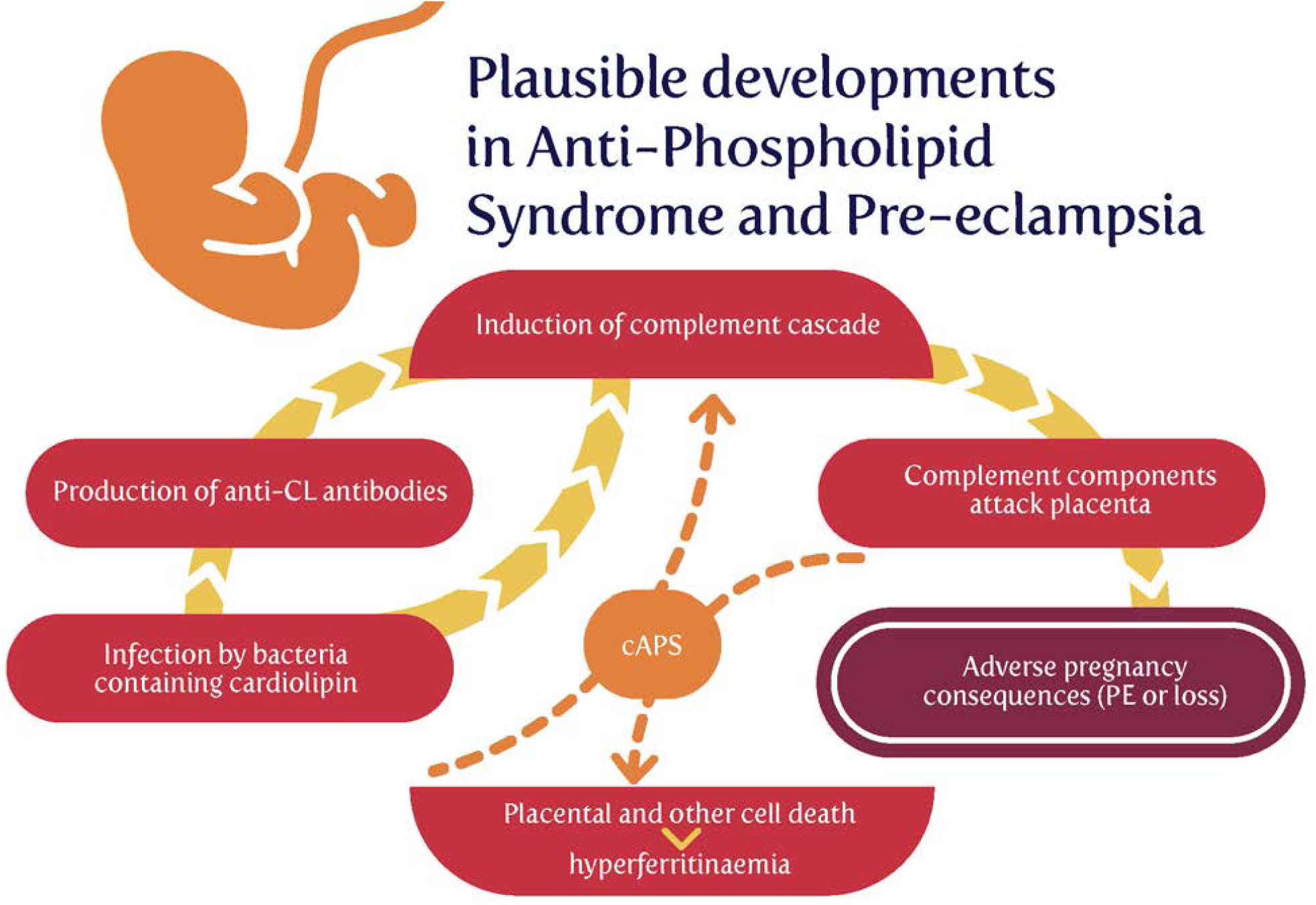
Possible relationships between cardiolipin exposure and disease sequelae.

## Treatment options based on (or consistent with) the ideas presented here

Although often unwritten or implicit, the purposes of much of fundamental biomedical science is to find better diagnostics and treatments for diseases (a combination sometimes referred to as theranostics). Consequently, our purposes here are to rehearse some of those areas where appropriate tests (in the form, ultimately, of randomised clinical trials, RCTs) may be performed. Clearly, as before [32], and recognising the issues of antimicrobial resistance, one avenue would exploit antibiotics much more commonly than now. We note that pharmaceutical drugs are prescribed or taken during 50% or more of pregnancies [769-778]. Anti-infectives are the most common such drugs, and some 20-25% of women or more are prescribed one or more antibiotics during their pregnancies [770; 771; 774; 776; 777; 779-782].

Given the role of male semen infection, we suggest that more common testing of semen for infection is warranted, especially using molecular tests. Our analyses suggest that antibiotics might also be of benefit to those males presenting with high microbial semen loads or poor fertility [783]. Another strategy might involve stimulating the production of antimicrobial peptides in semen.

Of the list of bacteria given in Table 2 as being associated with PE, *H. pylori* stands out as the most frequent. One may wonder why a vaccine against it has not been developed, but it seems to be less straightforward than for other infections [784; 785], probably because–consistent with its ability to persist within its hosts–it elicits only a poor immune response [786; 787]. Our own experience [788] is that many small molecules can improve the ability of other agents to increase the primary mechanisms that are the target assay, while having no direct effects on them themselves. Although ‘combinatorial’ strategies often lead to quite unexpected beneficial effects (e.g. [789; 790]), this ‘binary weapon’ strategy is both novel and untried.

As also rehearsed in more detail previously (e.g. [791; 792]) many polyphenolic antioxidants act through their ability to chelate unliganded iron, and thereby keep it from doing damage or acting as a source of iron for microbial proliferation. Such molecules may also be expected to be beneficial. Other strategies may be useful for inhibiting the downstream sequelae of latent infections, such as targeting inflammation or coagulopathies.

## Conclusions, summary and open questions

We consider that our previous review [32] made a very convincing case for the role of (mostly dormant) microbes in the aetiology of PE. However, we there paid relatively scant attention to two elements, viz (i) the importance of the immune system [145], especially in maternal immunotolerance, and (ii) the idea that possibly the commonest cause of the microbes providing the initial infection was actually infected semen from the father. We also recognise that epigenetic information [358; 793-795] can be provided by the father and this can be hard to discriminate from infection (if not measured), at least in the F_1_ generation. This said, microbiological testing of semen sems to be a key discriminator if applied. The ‘danger model’ [155; 157-160], in which it is recognised that immune activation owes more to the detection of specific damage signals than to ‘non-self’, thus seems to be highly relevant to PE [162].

Overall, we think the most important ideas and facts that we have rehearsed here include the following:

- Following Medawar’s recognition of the potential conundrum of paternal alloantigens in pregnancy, most thinking has focused on the role of maternal immunotolerance, and the role of regulatory T cells therein;
- Many examples show that sexual familiarity with the father helps protect against PE; however, this does not explain why in many cases exposure to paternal antigens is actually protective (and not even merely neutral);
- Semen contains many protective and immune-tolerance-inducing substances such as transforming growth factor β (TGF-β);
- However, semen is rarely sterile, and contains many microbes, some of which are not at all benign, and can be transferred to the mother during copulation;
- If one accepts that there is often a microbial component to the development of preeclampsia, and we and others have rehearsed the considerable evidence that it is so, then semen seems to a substantial, and previous largely unconsidered source of microbes;
- Some determinands, such as complement factor Bb, seem to reflect microbial infection and not just general inflammation that can have many other causes, and may therefore be of value in untangling the mechanisms involved;
- An improved understanding of the microbiology of semen, and the role of antibiotics and vaccination <u>in the father</u>, seems particularly worthwhile;
- Coagulopathies are a somewhat under-appreciated accompaniment to PE, and may contribute to its aetiology;
- The ‘danger model’ of immune response seems much better suited to describing events in pregnancy and PE than is the classical self/non-self analysis;
- The features of PE are not at all well recapitulated in animal models [24], and certainly not in rodents. However, it seems likely that they still have much to contribute [796; 797].

Open questions and further research agenda items include the following:

- There is a need for improved molecular and culture-based methods of detecting microbes in blood and tissues in which they are normally considered to be absent, both in the mother and the father;
- Notwithstanding the promise of metabolomics (see e.g. [798; 799]), there remains a need for better diagnostics, especially early in pregnancy;
- Issues of antimicrobial resistance are well known (e.g. [800-802]), and most antibiotics work only on growing cells, so there is a significant role for those that work on persisters and other non-replicating forms [803-805];
- The increasing online availability of patient information will permit greater exploitation to assess these ideas from an epidemiological point of view;

## Acknowledgements

DBK thanks the Biotechnology and Biological Sciences Research Council (grant BB/L025752/1) for financial support. LCK is a Science Foundation Ireland Principal Investigator (grant number 08/IN.1/B2083). LCK is also the Director of the Science Foundation Ireland-funded INFANT Research Centre (grant no. 12/RC/2272)

